# Transcriptomic profiling of the middle temporal gyrus reveals differential glial/neuronal dysregulation across Alzheimer’s disease and aging

**DOI:** 10.1101/2025.10.19.683343

**Authors:** IS Piras, A Bonfitto, S Song, A Aldabergenova, J Sloan, A Trejo, JC Troncoso, C Geula, EJ Rogalski, CH Kawas, M Corrada, TG Beach, GE Serrano, PF Worley, CA Barnes, MJ Huentelman

## Abstract

Alzheimer’s disease (AD), the most common cause of dementia, is characterized by amyloid-β plaques, neurofibrillary tangles, and widespread neuronal dysfunction. Aging, the strongest risk factor for AD, is also associated with some overlapping processes, such as neuronal cell transcriptional downregulation and glial cell activation. The middle temporal gyrus (MTG) is a brain region that supports semantic processing and default-mode connectivity and shows early vulnerability in both aging and AD. Here we profile bulk RNA-seq from 606 postmortem MTG samples with the goal of understanding the transcriptional changes associated with AD and aging. In 217 clinical and neuropathologically confirmed AD versus 290 no-dementia controls donors, we identify 613 differentially expressed genes (390 up, 223 down; |log2 fold change| ≥ 0.5; BH P < 0.05), with NPNT and ADAMTS2 among the top upregulated signals. Cell set enrichment indicates reduced excitatory neuronal signatures together with increased microglial, astrocytic, endothelial, and pericyte programs. Gene-set analyses reveal strong activation of angiogenesis, extracellular-matrix organization, wound response, adaptive immunity, and coordinated suppression of neuronal and mitochondrial processes, including synaptic signaling and respiratory-chain complexes. Multiscale coexpression mapping resolves three disease clusters: a neuron-mitochondrial module suppressed in AD (M5; hub PJA2; key driver GABRB3), a microglial immune module upregulated in AD (M6; hub C1QC; key driver FCER1G), and an increased astrocyte-vascular extracellular-matrix module in AD (M8; hub ESAM; key driver TAGLN). Across 324 non-AD controls aged 24–108 years, aging is associated with declines in gene expression associated with translation, proteostasis, and mitochondrial function and increases in those linked to oligodendrocyte and myelination programs (for example M4; hub CNTN2; key driver MOBP); in a 65+ subset, neuronal and protein-folding modules show the strongest decrements with reduced glial gene expression upregulatio. Our results indicate that late-life aging involves increased glial responses and neuronal/proteostasis suppression, whereas AD is also associated with immune– vascular–ECM activation and suppression of neuronal programs.

## Introduction

Alzheimer’s disease (AD) is the most common form of dementia and is clinically characterized by progressive cognitive decline and neuropathologically defined by amyloid-β plaques and neurofibrillary tangles^1,2^. Additionally, multiple cellular phenotypes have been reported, including synaptic and mitochondrial dysfunction, inflammation, vascular alterations, and impaired metabolic signaling^3–6^. The number of people with AD continues to rise globally, and aging remains the strongest risk factor^7^. Aging is characterized by overlapping, but not identical, transcriptomic changes such as downregulation of neuronal and mitochondrial pathways and increased glial gene expression^8,9^.

High-throughput transcriptomic profiling has been relevant for the characterization of these processes in AD and RNA-seq datasets have highlighted the down-regulation of synaptic and mitochondrial genes with concomitant up-regulation of immune and vascular pathways^10–19^. However, the middle temporal gyrus (MTG) has not been extensively characterized with RNA-sequencing^20^. The MTG is a brain region that supports lexical-semantic processing and conceptual knowledge critical for language comprehension^21,22^. The MTG also participates in the default mode network^23^, the activity of which was linked to differences between healthy aging and AD over 20 years ago^24^. With typical aging, cortical thinning consistently affects the temporal gyri of the brain, including the MTG, and this has been associated with declines in higher-order cognition^25^. In AD, the MTG shows early atrophy and hypometabolism, and degeneration in this region can predict progression from typical aging to AD^26,27^. These findings suggest that the MTG is a key region for semantic cognition whose vulnerability is associated with both cognitive aging and AD-related pathology.

In this study, we performed an extensive transcriptomic analysis of more than 600 postmortem samples from the MTG, encompassing individuals with clinical and neuropathologically confirmed AD and no-dementia controls (CTL). We also conducted a detailed investigation of aging in individuals without dementia and significant AD pathology. This large dataset allowed us to examine both AD-associated and aging-related transcriptional changes within the same cortical region. By combining differential gene expression with coexpression network analyses, we mapped the transcriptional programs associated with AD pathology and brain aging. This combined strategy allowed us to identify molecular changes that are shared between aging and AD from those that are unique to each condition. We provided insights into the distinct and overlapping mechanisms driving age-and AD-related transcriptional remodeling in the human brain.

## Methods

### Samples

Fresh frozen cortical sections from the middle temporal lobe were obtained from four different brain banks: Banner Sun Health Research Institute, the 90+ Study of the University of California Irvine, Johns Hopkins University, the SuperAging Research Initiative and Northwestern University brain bank. After quality controls (sequencing quality and outliers’ removal, see results section for details), we included a total of 606 samples. Neuropathological and demographic variables for the final samples included in the study are reported in **Table 1**. Informed written consent was obtained from all participants prior to inclusion in the study. There were two primary studies: 1) the AD study, which focused on identifying transcriptional signatures and co-expression networks associated with AD, and 2) the aging study, which examined transcriptional signatures across aging in no-dementia individuals.

**Table 1.** Demographic and neuropathological characteristics of the analyzed samples.

| Group | AD (217) | CTL (290) | *CTL (all ages) (n = 324) | **CTL (65+) (299) |
| --- | --- | --- | --- | --- |
| Age (average- median - range) | 91.8 \| 94.5 \| 65 - 110.6 | 92.2 \| 94.5 \| 65 - 108.6 | 88.7 \| 93 \| 24 - 108.6 | 92.2 \| 94 \| 65 \| 108.6 |
| Sex ratio (F:M) | 2.14 | 1.29 | 1.23 | 1.27 |
| Braak Stage (0 \| 1 \| 2 \| 3) | 0\|7\|64\|146 | 0\|52\|195\|43 | 4\|65\|195\|43 | 0 \| 52 \| 195 \| 43 |
| CERAD plaque score (0 \| 1 \| 2 \| 3) | 13\|13\|32\|159 | 78\|58\|49\|105 | 94 \| 59 \| 49 \| 105 | 78\|58\|49\|105 |
| PMI (hours) (average- median - range) | 4.04 \| 3.62 \| 1.5 - 18.5 | 4.36 \| 3.88 \| 1 - 20.8 | 4.97 \| 3.91 \| 1 - 46 | 4.70 \| 3.90 \| 1 - 46 |

\* Neuropathological data non available for 17 donors
\*\* Neuropathological data non available for 9 donors

For the AD study, we included individuals with a clinical and neuropathological diagnosis of AD dementia and no-dementia controls (CTL) with age at death equal to or greater than 65 years old. All AD cases selected had a clinical diagnosis of AD dementia , Braak Stage III-VI, and/or CERAD plaque density “moderate” or “frequent” (AD high pathology). Using these criteria, we selected a total of 217 AD and 290 CTL participants. For the aging study, we included all the no-dementia controls (CTL) without age cutoff, plus 9 participants from the SuperAging research Initiative^28–31^, not pathologically confirmed but classified as “superagers” using cognitive tests. The final sample size for the aging CTL group was 324. This study samples were further stratified into “all-ages” (24-108 years; n = 324) and “65+”, including in the latter only donors aged 65 or older (n = 299).

### Brain Bank Contributors

#### Banner Sun Health Brain and Body Donation Program (BBDP)

The Arizona Study of Aging and Neurodegenerative Disorders and Body Donation Program at Banner Sun Health Research Institute is a longitudinal clinicopathologic cohort initiated in 1987 that recruits mainly from retirement communities in metropolitan Phoenix, Arizona^32,33^. Enrollees undergo standardized medical, neurologic, and neuropsychological assessments during life, with rapid autopsy at death (median post-mortem interval ∼3.0 hours) yielding high-quality frozen tissues, including a median brain RNA Integrity Number of 8.9^32,33^. Whole-body donation has been available since 2005, and both anatomical and neuropathologic diagnoses are rendered by licensed pathologists using contemporary consensus criteria^33^. A total of 232 samples were included in the AD/CTL study, and 151 in the aging study.

#### 90+ Study

The 90+ Study is a community-based longitudinal cohort of adults aged 90 years and older, drawn primarily from surviving members of the Leisure World Cohort in Laguna Woods, California^34^. Participants are evaluated approximately every 6 months with a neurological examination, a standardized neuropsychological battery, and informant questionnaires; remote phone or mail assessments are used when in-person visits are not feasible^34,35^. Cognitive status (normal cognition, CIND, or dementia) was assigned after a participant’s death in a multidisciplinary diagnostic conference that used information from all longitudinal evaluations, informants, laboratory tests, and medical records, with conferences blinded to pathological evaluation.^30^ Neuropathological evaluation was done blinded to clinical information using the National Institute on Aging-Alzheimer’s Association “ABC” score^36,37^. ^34^Additional cohort descriptions and epidemiologic findings on dementia prevalence and incidence in this population have been published^38,39^. A total of 275 samples were included in the AD/CTL study and 156 in the aging study.

#### SuperAging Research Initiative

This longitudinal cohort enrolls community-dwelling adults aged 80 years or older who perform at or above the mean for 50–60-year-olds on Rey Auditory Verbal Learning Test (RAVLT-DR) delayed recall and at least within the average range for their age on non-memory measures; recruitment occurs through the Alzheimer’s Disease Center and community outreach^40,41^. It also enrolls cognitively average controls who perform within the average normative range on the RAVLT-DR for their demographics and at least within the average range for their age on non-memory measures. Participants complete standardized neuropsychological evaluations at baseline and follow-up, with optional neuroimaging and biomarker studies and brain donation for neuropathology^41^. This study was expanded in 2021 as a multisite with enrollment across five sites in the US and Canada^28^. A total of 9 samples were included in the aging study.

#### Johns Hopkins Division of Neuropathology Brain Bank

Tissues were fresh frozen punches of the left MTG. This autopsy cohort draws brain donations through the Maryland Office of the Chief Medical Examiner and affiliated programs, explicitly including young adults; standardized dissections and region-specific sampling are performed at the Johns Hopkins Division of Neuropathology^42,43^. Each case, detailed medical and toxicology histories are abstracted, and tissue is processed for uniform histology, with genotyping and molecular assays performed when applicable^42,43^. The cohort has been used to characterize preclinical Alzheimer-type pathology in individuals aged 40–50, informing age-stratified analyses of early disease biology^43^. A total of 8 samples were included in the aging study.

### Data analysis

FASTQ files were analyzed using the Nextflow RNA-seq pipeline, including trimming with Trim Galore, alignment with STAR^44^, and quantification of counts at the gene level with SALMON. Quality control was conducted using MultiQC and Qualimap^45^. Samples were included if they had all covariates (age, sex, RIN, and PMI), at least 20 million sequencing reads, and if 80% of those reads were uniquely mapped to the human transcriptome. Raw counts were imported from DESeq2^46^ and, for quality control purposes, transformed using the *vst* method. We removed genes located on the sex chromosomes, and conducted Principal Component Analysis (PCA) to identify outlier samples, defined as those that were above ± 3 standard deviations from the average of at least one of the two top PCs. The relationship between gene expression and potential confounding variables (RNA integrity number RIN, postmortem interval PMI, sample source, sex and age) was investigated by correlating the top two PCs with each of the potential confounding variables and assessing significance with Pearson’s r (for RIN, age, and PMI) or Wilcoxon Test (source and sex). For normalization and differential expression, we generated three different datasets with DESeq2: AD/CTL, aging-65+ ( ≥ 65 years old) and all-ages. This approach was implemented to minimize bias by excluding samples irrelevant to each specific analysis, thereby reducing their influence on the overall variance. Each dataset was prefiltered for low expressed genes using a minimum count threshold equal to the total sample size. Normalization and differential expression were conducted using DESeq2^46^, including all of the covariates in the model (age, sex, PMI, RIN, source and sequencing batch). The continuous covariates were standardized using the ‘*scale’* function from R, while differential expression p-values were adjusted for multiple testing using the Benjamini & Hochberg (BH) method^47^. Log2 Fold Change (LFC) was estimated using the shrinkage method (*apeglm*)^48^ to avoid inflated LFC and p-values due to transcripts with very low counts (these were found to mostly be non-protein coding transcripts). During differential expression, we applied the independent filtering method implemented in DESeq2 to remove lowly expressed genes, optimizing the cutoff for α = 0.05. Genes with BH adjusted p-values (BHP) < 0.05 and absolute log_2_FC (LFC) ≥ 0.25 were considered statistically significant. Boxplots (AD/CTL study) and scatterplots (aging study) were visualized using variance-stabilizing transformed data, adjusted for covariates. The adjustment was conducted using the *removeBatchEffect* function for the R-package *limma*^49^.

All lists of genes were further analyzed using the Gene Set Enrichment Analysis (GSEA) referenced to the Gene Ontology database, as implemented in the R-package *clusterProfiler*^50^. Redundant GO functional classes were removed using the ‘*simplify*’ function with the default settings. GO processes with BHP < 0.05 were considered statistically significant. Cell-set enrichment analysis (CSEA) was conducted using as gene sets the lists of cell-specific markers from the single nucleus RNA sequencing study from Mathys et al.^18^, obtained using our deconvolution method as previously described^51^. The enrichment analysis was conducted using the fast enrichment analysis (fGSEA) method^52^.

Coexpression analysis was conducted using the Multiscale Embedded Gene Expression Network Analysis (MEGENA) algorithm^53^ only selecting the protein coding genes after annotation with *Biomart*. As input, we used the data matrices previously generated for the boxplot and scatterplot visualizations (see above). The datasets were further filtered to include only the top 50% of genes with the highest median absolute deviation. For coexpression module generation, we first calculated signed pairwise gene correlations using Pearson’s method with 1,000 permutations, retaining correlations that were significant at the 5% FDR level (function: calculate.correlation). Significantly correlated gene pairs (FDR < 0.05) were ranked and iteratively tested for planarity, leading to the development of a planar filtered network using the planar maximally filtered graph technique (function: calculate.PFN). Subsequently, we conducted a multiscale clustering analysis to identify coexpression modules at varying network scale topologies and their respective hub genes (function: do.MEGENA). Coexpression modules deemed significant (with a permuted P < 0.01 and module of 50 genes or more) were carried forward for further analysis. Next, we extracted module eigengenes (the first principal component obtained from the module genes) using the function moduleEigengenes from the WGCNA R package^54^. Pairwise differential expression between diagnostic groups (AD vs CTL) and across aging in CTL subjects was computed using the *limma* R-package. Modules with significant associations were annotated for GO functional classes using a hypergeometric testing as implemented in *clusterProfiler*. Additionally, we investigated the cell-specific enrichment of each associated modules by hypergeometric statistics (*bc3net* R-package) using brain-specific cell markers as described in our previous study^51^. Weighted Key Driver Analysis (wKDA) was conducted using the Mergeomics webtool^55^, applying the following parameters: search depth of 1, edge type undirected, min hub overlaps 0.33 and edge factor 0. We used the available Bayesians networks from cortex (GTEx v8) and the ‘legacy’ brain network.

### Large Language Model Use

A large language model (LLM) was used during manuscript preparation to check spelling and grammar, improve readability, and broaden the manuscript’s accessibility to scientific disciplines beyond those specializing in the article’s main topics.

## Results

### Filtering and quality controls

We sequenced a total of 621 samples, with an average number of 31.8 million reads (range: 16.7 million – 52.2 million; SD: 5.8 million) (**Fig. S1**). We removed two samples with fewer than 20 million reads. The average mapping rate, as determined by STAR (uniquely mapped reads), was 93.7% (range: 67.9% - 95.8%; SD: 1.85) (**Fig. S2**). We removed two samples with a mapping rate below 80%, obtaining a final sample size of 617. Data were prefiltered to remove lowly expressed genes (genes with total counts across all samples < 617), resulting in 38,603 genes. Then, counts were transformed using the variance-stabilizing transformation (VST) method, and we extracted the top two principal components, classifying 11 samples as outliers (**Figs S3** and **S4**). After we removed the outliers, the final dataset consisted of 606 samples. We explored the relationship between gene expression and potential confounding factors that correlated with the top two principal components, detecting a significant association of one of the two components with the following variables: age, postmortem interval (PMI), RNA integrity number (RIN), sample source (brain bank), and sex (**Table S1**). Among the variables tested, only the sequencing run demonstrated a lack of significant association with gene expression (P > 0.221). All of these confounding variables were included in the downstream analyses as covariates and were adjusted for in the model.

### Transcriptomic changes in the AD MTG are associated with down-regulation of synaptic and mitochondrial pathways as well up-regulation of immune, vascular and extracellular matrix processes

For the AD study, we included samples that met the criteria described in the Methods section and excluded those with missing covariates or neuropathological variables, obtaining a final sample size of 217 AD and 290 CTL participants. Differential expression analysis identified a total of 613 significant genes (LFC ≥ 0.5 and BHP < 0.05), with 390 overexpressed and 223 underexpressed in AD (**Fig. 1A, Table S2**). Among the top protein-coding genes, we identified *NPNT*, *ADAMTS2*, *GFAP* and *ECM2* (**Fig. 1B**). CSEA analysis revealed a significant enrichment of excitatory neuronal genes (underexpressed in AD), along with microglia, astrocyte, endothelial, and pericyte genes (overexpressed in AD) (**Fig 1C**; **Table 3S)**. GSEA analysis revelated a strong upregulation of processes associated with angiogenesis, extracellular matrix organization, wound response, and adaptive immune activation (NES > 2.0; BHP < 1.0 × 10^-12^). In contrast, neuronal and mitochondrial processes, including synaptic signaling, synaptic membrane organization, and respiratory chain/ATP synthesis complexes, were significantly underexpressed in AD (NES < -2.0, BHP = 1.0 × 10^-06^) (**Figs. 1D** - **1F, Table S4**). We then constructed a multiscale coexpression network using only protein coding genes and, after eigenvalue extraction, identified 61 modules significantly associated with AD (**Fig 1G**, **Table S5;** see **Table S6** for the module specific GO functional classes; see **Table S7** for the list of significant key drivers). These modules were grouped into three main functionally distinct clusters, including the top modules M5, M6 and M8 (**Fig. 1H**). The largest cluster included module M5 (n = 3,786; hub gene: *PJA2*) as a top-level module, underexpressed in AD and enriched for ribosomal and mitochondrial functions and showing additional signals from synaptic and ion transport processes (**Fig. 1I**). Additionally, module M5 was particularly expressed in excitatory neurons (BHP = 7.8 × 10^-64^) with some signal in inhibitory neurons (BHP = 3.1 × 10^-06^). The top key driver of this module was *GABRB3* (FDR = 6.4 × 10^-08^), also showing an under-expression trend in AD (LFC = -0.080; P = 1.5 × 10^-02^; BHP = 1.0 × 10^-01^). The second cluster included module M6 (n = 635; hub gene: *C1QC*), overexpressed in AD and enriched for immune system-related GO terms and microglia-expressed genes (BHP = 2.0 × 10^-242^) (**Fig. 1L**). The top key driver was *FCER1G* (FDR = 5.1 × 10^-32^), also upregulated in AD (LFC = 0.148; P = BHP = 6.6 × 10-^02^). The third cluster included module M8 (n = 758; hub gene: *ESAM*), overexpressed in AD, enriched for extracellular matrix organization and blood vessel development. This module was primarily associated with astrocyte-expressed genes (p = 2.2 × 10^-66^), with *TAGLN* identified as the top key driver included in the module (FDR = 2.1 × 10^-20^). This gene showed a non-significant upregulation in AD (LFC = 0.036; p = 1.7 × 10-01; BHP = 4.6 × 10^-01^).

**Figure 1.**
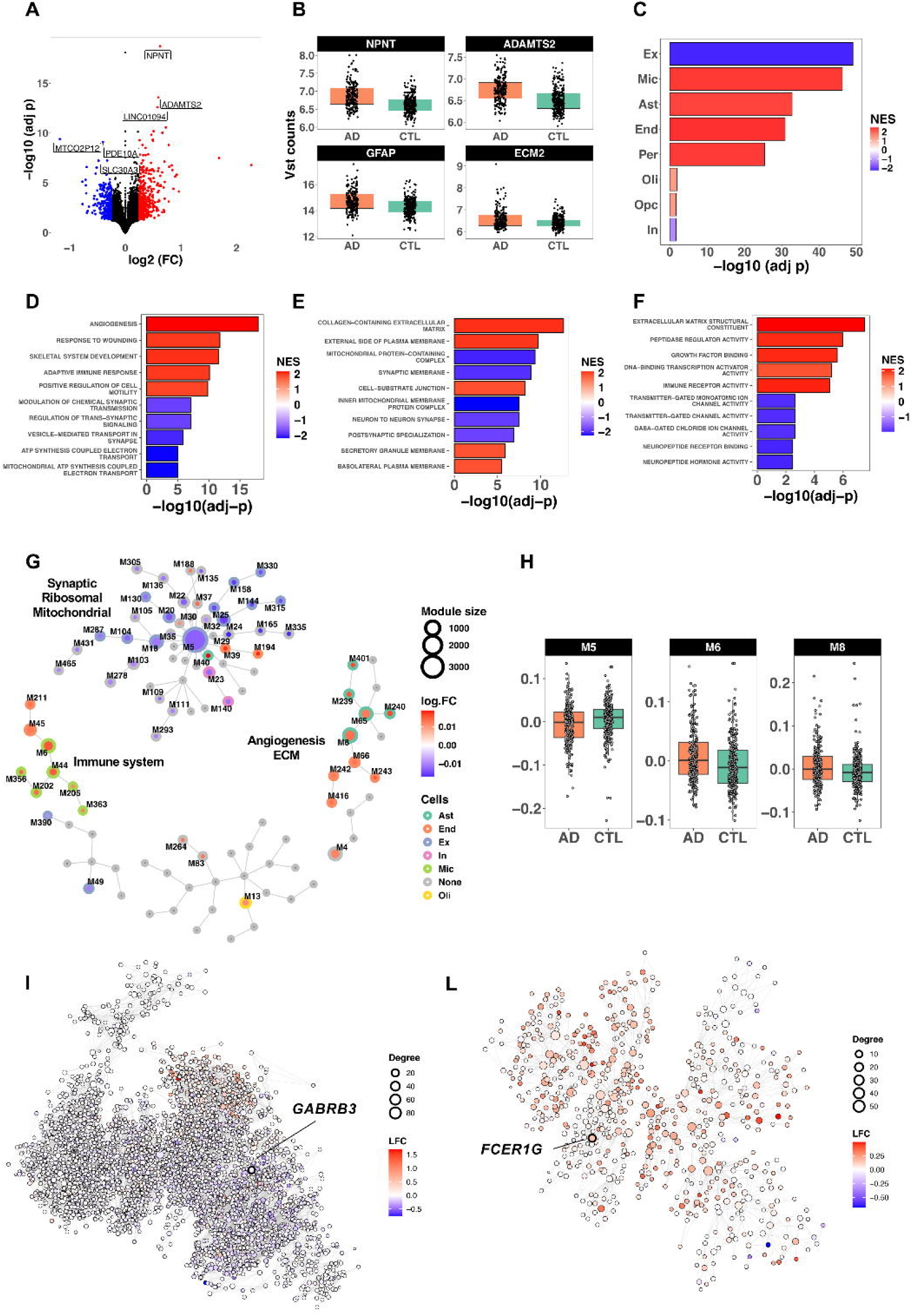
Transcriptomic alterations in the middle temporal gyrus (MTG) of AD compared to control (CTL) brains. (A) Volcano plot of differential expression analysis. Genes significantly overexpressed and underexpressed in AD are shown in red and blue, respectively (n = 613 DEGs, BH-adjusted *p* < 0.05, |log_₂_FC| ≥ 0.25). (B) Top four protein-coding differentially expressed genes identified. (C) Cell set enrichment analysis showing reduced expression of excitatory neuronal genes (Ex) and increased expression of microglial (Mic), astrocytic (Ast), endothelial *End), and pericyte-associated genes in AD. (D–F) Gene set enrichment analysis (GSEA) results using Gene Ontology for (D) Biological Processes, (E) Cellular Components, and (F) Molecular Functions. (G) Overview and relationships of coexpression modules identified in MTG. Each node represents a coexpression module; the border color indicates cell-specific enrichment, while the core reflects the log_2_FC of the module eigenvector differential expression between AD and CTL. (H) Differential expression of top-level, functionally relevant coexpression modules associated with AD compared to CTL. (I) Synaptic/mitochondrial/ribosomal module Mt5 coexpression network; node colors represent differential expression (log_₂_FC) in AD vs. CTL. The top key driver gene (*GABRB3*) is indicated. (L) Immune-related module M6 coexpression network; node colors represent log_₂_FC direction in AD vs. CTL. The top key driver gene (*FCER1G*) is indicated.

### Aging stages are characterized by differential cell-specific expression patterns

For the aging study, we explored the relationship between gene expression and aging in a sample of 324 non-AD controls aged between 24 to 108 years (referred to as the “all-ages” group), followed by an analysis of donors aged 65 years and older (“65+” group; n = 299). The goal was to identify aging-stage-specific expression patterns. We identified 47 genes significantly associated with chronological age in the all-ages group (LFC ≥ 0.25; BHP < 0.05), 25 of which were downregulated (**Fig. 2A**, **Table S8A**). The same analysis conducted in the 65+ group revealed only nine differentially expressed genes, most of which were downregulated (**Table S8B**). Only two genes overlapped between the two groups: *GPR26* and *NEB* **(Fig 2B)**. The top protein-coding genes significantly associated in the all-ages group were: *EDN3*, *TTR*, *TNFRSF19* and *PRLR*, whereas in the 65+ group, we identified *GPR26*, *FGF18*, *HSPA6* and *NEB.* CSEA analysis **(Fig 1C)** in the all-ages group confirmed the downregulation of excitatory neuronal genes and revealed increased expression of astrocyte- and oligodendrocyte-associated genes (**Table S9A**). In contrast, the 65+ group showed a stronger neuronal signature, with significant downregulation of both excitatory and inhibitory neuronal genes associated with increased age, with minimal glial-related gene upregulation (**Table S9B**). GO-GSEA analysis in the all-ages group indicated strong downregulation of pathways related to ribosomes, protein-folding, proteostasis, mitochondria, and hormone signaling, suggesting impaired translation and proteostasis in this group. The only positively enriched process among the 74 significant categories was “intracellular chloride channel activity” (**Table S10; Figs D-E**). In the 65+ group, the same analysis revealed a large proportion of functional categories with negative enrichment scores (**Table S11**). The strongest decreases were linked to “blood microparticle” and proteostasis/heat-shock GO classes, including unfolded-protein binding and protein-folding chaperones. Additionally, neuronal/synaptic functional classes were also underexepressed with aging. Only two categories were overexpressed: “regulation of cell communication by electrical coupling” and “proximal/distal pattern” formation.

**Figure 2.**
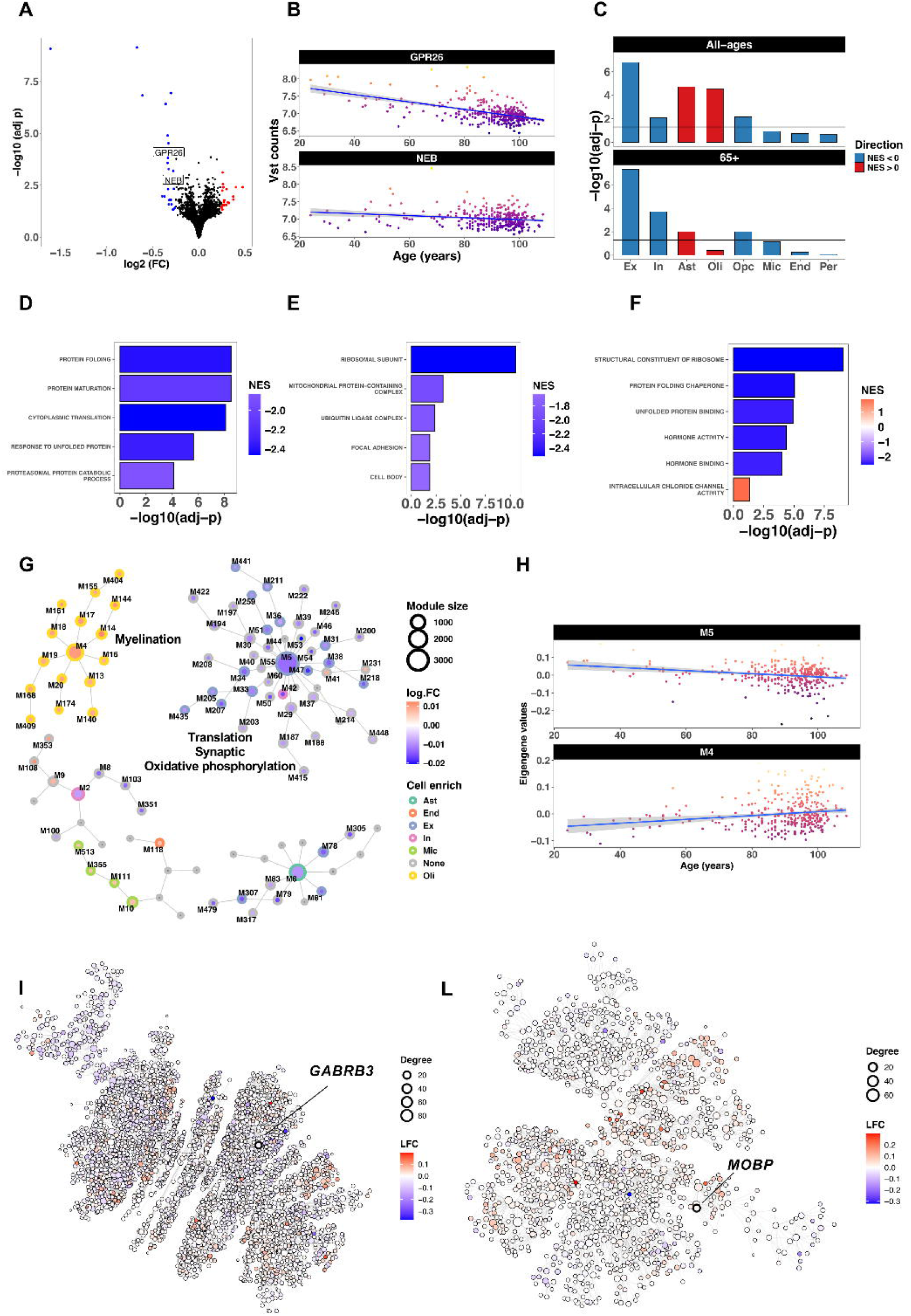
Transcriptomic alterations in the middle temporal gyrus (MTG) of aging. (A) Volcano plot of differential expression analysis in the all-ages cohort (24-108). Genes significantly overexpressed and underexpressed with aging are shown in red and blue, respectively (n = 47 DEGs, BHP < 0.05, |log_₂_FC| ≥ 0.25). Labels indicate the two genes that also overlapped with the 65+ group. (B) Genes significant in both all-ages and 65+ groups. (C) Cell set enrichment analysis showing reduced neuronal gene expression in the 65+ group and an increase of glial signatures in the all-ages group. (D–F) Gene set enrichment analysis (GSEA) results using Gene Ontology for (D) Biological Processes, (E) Cellular Components, and (F) Molecular Functions. (G) Overview and hierarchical relationships of coexpression modules identified in the all-ages group. (H) Differential expression of the two top-level, functionally relevant coexpression modules associated with aging in the all-ages group. (I) Translation/synaptic/oxidative phosphorylation module M5 coexpression network; node colors represent differential expression across aging. The top key driver gene (*GABRB3*) is indicated. (M) Myelination/oligodendrocyte-related module M4 coexpression network; node colors represent log_₂_FC direction in aging. The top key driver gene (*MOBP*) is indicated.

MEGENA analysis in the all-ages group identified age-related modules, which we grouped into two main clusters (**Fig. 2G** and **Fig 2H**, **Table S12**; see **Table S13** for the GO classes associated with modules; see **Table S14** for the list of significant key drivers). The main cluster was centered around module M5 (n = 3,418; hub gene *PJA2*), which was downregulated with aging and enriched for excitatory neuron expression and translation/ribosome functional classes (**Fig. 2I**). The key driver was *GABRB3* (FDR = 7.4 × 10^-12^), which showed a non-significant downregulation trend with age (LFC = 0.018; P = 2.1 × 10^-01^). The M5-related modules were associated with synaptic signaling and oxidative phosphorylation and were enriched for excitatory neuron-expressed genes. The second largest cluster was centered around the top-level module M4 (n = 1,326; hub gene: *CNTN2*), which was upregulated with aging and enriched for myelination processes and oligodendrocyte signatures (**Fig. 2L**). The top key driver of M4 was *MOBP* (FDR = 9.3 × 10^-35^), which also showed a non-significant upregulation trend with age (LFC = 0.012; P = 1.1 × 10^-01^). All M4 related modules were enriched for oligodendrocyte genes, and the largest of these (M14; n = 217) was associated with myelin sheath. The same analysis conducted in the 65+ aging group identified a total of 17 modules associated with age (**Table S15;** see **Table S16** for the GO classes associated with modules; see **Table S17** for the list of significant key drivers). The largest group of related modules included M2 (n = 599, hub gene: *HGSNAT),* which was underexpressed with aging and enriched for protein-folding GO functional classes. The second group included M112 (n = 298; hub gene: *SULT4A1*), which was downregulated with age and enriched for synaptic processes as well as for inhibitory neuron genes.

## Discussion

In this study, we characterized MTG from a large sample of postmortem brains, first investigating AD signatures and then exploring their relationship with normal aging (range: 24-108 years old). We analyzed 217 AD and 290 CTL brains, detecting 613 DEGs, the majority of which (63.6%) were overexpressed in AD. The top gene identified was *NPNT* (LFC = 0.625; BHP = 1.9 × 10^-19^). *NPNT* (nephronectin) encodes an extracellular-matrix glycoprotein containing and RGD integrin-binding motif^56^ and plays a known role in matrix organization and cell adhesion. Notably, this gene was recently reported as the top differentially expressed gene in a cross-ancestry transcriptomic meta-analysis^57^. The second most significant gene was *ADAMTS2* (LFC = 0.594; BHP = 2.7 × 10^-14^) (ADAM metallopeptidase with thrombospondin type 1 motif 2), which encodes a zinc metalloprotease. In agreement with our results, increased expression of this gene has been associated with cognitive decline in the anterior cingulate cortex and reported in a multiethnic transcriptome AD meta-analysis^58,59^. The other top genes, ECM2 and GFAP, were associated with increased ECM signatures and astrocyte activity, respectively. Overall, our differential expression results revealed a stronger deregulation of excitatory than inhibitory, neuronal genes. Accordingly, we observed suppression of synaptic and mitochondrial respiratory/ATP synthases functional classes, suggesting neurodegeneration, impaired neurotransmission and disrupted bioenergetic mechanisms. Conversely, we found a larger number of overexpressed non-neuronal genes in AD, primarily associated with microglia, astrocyte, endothelial cells and pericytes. Consistently, we observed increased immune, vascular and ECM transcriptional programs linked to tissue repair/remodeling and neuroinflammation. These phenomena have also been reported in previous bulk tissue and single-nuclei RNA-seq data^13,18,60–62^. Changes in astrocyte and ECM upregulated in AD are compatible with synaptic dysfunction. Astrocytes reside in the neurovascular-synaptic interface, and the ECM regulates synaptic integrity and plasticity. Our data suggest an excess of ECM accumulation, which may constrain activity dependent remodeling and microglial pruning that support cognition^63,64^. Unlike normative aging, AD shows coordinated immune-vascular ECM activation along with suppression of neural and synaptic programs.

Our multiscale coexpression analysis recapitulate these molecular features in AD. Synaptic and mitochondrial suppression were detected in the large module M5, which was downregulated in AD. The key driver of this module was *GABRB3*, encoding for the β3 subunit of the GABA-A receptor, which showed a downregulation trend in AD. The gene has been consistently reported as downregulated in AD^65^. The increase in immune processes was reflected in M6-related modules, which were upregulated and had *C1QC* as a hub gene. The key driver of this module was *FCER1G* (FDR = 5.1 × 10^-32^), upregulated in AD (LFC = 0.144; BHP = 6.6 × 10^-02^). *C1QC* encodes the C1q subcomponent subunit C, part of the C1q complex, which is the recognition component of the classical complement pathway, associated with the downstream inflammatory response. The C1q complex interacts with TREM2^66^ and has been implicated in synapse elimination mediated by astrocyte and microglia in AD^67^. Additionally, it enhances microglia activation through a positive feedback loop, increasing neuroinflammation^68,69^. *FCER1G,* which encodes the Fc receptor γ chain, recruits Spleen Tyrosine Kinase to activate key innate immune effector functions such as phagocytosis and cytokine production. It has been described as hub-gene in microglia enriched modules associated with aging and neurodegeneration, including AD^70^. Finally, module M8 summarized the endothelial/ECM component. This module was upregulated in AD, with *ESAM* as a hub gene (significantly upregulated in AD: LFC = 0.229; BHP = 6.9 × 10^-04^), enriched for astrocyte genes. The top key driver was *TAGLN* (FDR = 1.5 × 10^-10^), which showed a slight increase in AD. *ESAM* (Endothelial Cell Adhesion Molecule) encodes a transmembrane immunoglobulinl1llike adhesion protein located at endothelial tight junctions and expressed on platelets. Its function is related to maintaining endothelial barrier integrity and regulating leukocyte transmigration, as well as playing a role in angiogenesis^71^. This gene may be involved in the vasculature remodeling and maintenance of the blood brain barrier following AD-related injury.

The aging study highlighted two genes *GPR26* (G protein–coupled receptor 26) and *NEB* (nebulin) – that were decreased in expression with advancing age in both the all-ages (24-108 years old) and 65+ groups. *GPR26*, is primarily expressed in the brain and has been implicated in the maintenance of neuronal excitability and energy balance. NEB, a large actin-binding protein, might be involved in broader cytoskeletal remodeling processes. Notably, we identified distinct patterns at the cellular level across different age stages. In the all-ages group, the major shift in transcriptomic dysregulation appears to be associated with a decrease of excitatory neuronal gene expression and an increase in astrocyte and oligodendrocyte gene expression. These patterns, characterized by reduced neuronal and decreased glial increased expression, have been previously reported in human transcriptomics aging studies^9^. In the all-ages group, GSEA analysis revealed strong downregulation in pathways related to ribosomes, protein-folding, proteostasis, mitochondrial, and hormones, highlighting impaired translation and proteostasis^72^. These processes, especially mitochondrial and ribosomal functions, and the decreased expression in excitatory neuronal regions were summarized by the module M5, which included *GABRB3* as a key driver gene, showing a downregulation trend in aging. Notably, we identified a similar large module with *GABRB3* as a key driver, suggesting shared mechanisms of vulnerability between AD and aging. In addition, coexpression analysis highlighted a set of modules upregulated across aging and enriched for oligodendrocyte genes. The top module among these was M4 (key driver: *MOBP*), which showed an upregulation trend with age. This result suggests active compensatory remodeling of the myelin in the aging brain. MOBP is a myelin-specific structural protein mostly expressed in oligodendrocytes, playing a key role in myelin stabilization and integrity of myelin in the central nervous system^73^. This gene has been reported to be a key gene downregulated in Multiple System Atrophy^74^. Among the top DEGs we identified *EDN3* (Endothelin-3) and *TTR* (Transthyretin), both of which were downregulated with aging. *EDN3* is a vasoactive peptide crucial for vascular function and cardiovascular aging, although its role in brain aging remains unknown. *TTR* transports thyroxine and retinol in the cerebrospinal fluid and plasma. Its levels decline with age, likely due to the deterioration of the choroid plexus integrity and disrupted hormone transport^75^. Other studies suggest that lower circulating TTR levels are associated with all-cause mortality and risk of heart failure, making it a robust biomarker of systematic aging^76^.

In the 65+ group, transcriptomic dysregulation was much lower, with only nine genes significantly associated with aging, eight of which were downregulated. Similarly to the all-ages groups, we observed a decrease of both inhibitory and excitatory neuron gene expression, with lower changes in astrocyte and not significant enrichment for oligodendrocyte gene expression. This was confirmed by GO analysis, which showed downregulation of neuronal and synaptic processes, in addition to proteostasis and heat-shock functional classes, and in the coexpression analysis, which demonstrated an absence of glial-driven modules. These finding suggest that glial gene expression changes are more prominent in early aging than in later life, where neuronal gene expression downregulation is still the dominant pattern. However, these results should be interpreted with caution, as the small proportion of donors under 65 years old (7.7% of the sample) limits statistical power and highlights the need for confirmation in larger, more age-balanced cohorts.

It is important to note several potential limitations of our work. We acknowledge the differences between brain bank sources (e.g., characteristics of participants, staining methods, tissue sampling and assessment methods) and the possibility that those may influence the statistical outcomes of our analyses. We attempted to address this by investigating key variables that could be confounding, such as brain bank, RIN, PMI, sequencing run, age at death and sex, and by including those variables that were significantly correlated with the top two principal components in our statistical modeling. Secondly, we were able to include MTG specimens from donors spanning eight decades of age, however, brains from individuals under the age of 65 were much less prevalent in our study cohort. Therefore, some of the age-related findings that we describe could be artificially enhanced by the smaller number of cases examined from younger ages. We divided our cohort into AD cases and controls wherein the control samples were from individuals who did not have a dementia diagnosis while living, however, it is important to remember that cognitive performance is a continuum and our control cohort included individuals with mild cognitive impairment as well as individuals who had much higher than typical cognitive performance, such as the SuperAger donors from the Northwestern brain bank. Lastly, bulk-based transcriptome analysis is unable to fully discern cell-specific alterations and therefore our findings should be considered a starting point for understanding the changes happening in the MTG during aging and AD. Future work should include the use of single cell/nucleus RNA sequencing as well as spatial transcriptomic investigations, both of which will serve to further add additional information to what we report on here using bulk RNA sequencing.

In conclusion, our analysis of post-mortem MTG reveals a broad downregulation of excitatory neuronal, synaptic and mitochondrial processes, along with an upregulation of immune, vascular, and ECM pathways. The results are mostly confirmatory of prior published work, but no other in-depth analyses have been conducted on the MTG using high-throughput RNA sequencing. Additionally, we highlighted a few key genes associated with large coexpression networks perturbed in AD, such as *GABRB3* and *FCER1G*. The aging study revealed partly overlapping features, with more pronounced glial activation in early stages, driven by *MOBP*, followed by neuronal deregulation in later stages, driven by *GABRB3*, the same key driver identified in a large AD coexpression network. Finally, we highlighted the immune and vascular transcriptional programs in AD, which are not found to be significantly associated with aging in general in our dataset.

## References

1. Braak H, Braak E. Neuropathological stageing of Alzheimer-related changes. Acta Neuropathol. 1991;82(4):239–259. PMID: 1759558

2. 2024 Alzheimer’s disease facts and figures. Alzheimers Dement 2024 May;20(5):3708–3821. PMID: 38689398

3. Scheffer S, Hermkens DMA, Van Der Weerd L, De Vries HE, Daemen MJAP. Vascular Hypothesis of Alzheimer Disease: Topical Review of Mouse Models. Arterioscler Thromb Vasc Biol; 2021 Apr 1;41(4):1265–1283. PMID: 33626911

4. Cai Q, Tammineni P. Mitochondrial Aspects of Synaptic Dysfunction in Alzheimer’s Disease. Journal of Alzheimer’s Disease. IOS Press; 2017. p. 1087–1103. PMID: 27767992

5. Zhang J, Zhang Y, Wang J, Xia Y, Zhang J, Chen L. Recent advances in Alzheimer’s disease: Mechanisms, clinical trials and new drug development strategies. Signal Transduction and Targeted Therapy. Springer Nature; 2024. PMID: 39174535

6. Breijyeh Z, Karaman R. Comprehensive Review on Alzheimer’s Disease: Causes and Treatment. Molecules. MDPI; 2020. PMID: 33302541

7. Xiaopeng Z, Jing Y, Xia L, Xingsheng W, Juan D, Yan L, Baoshan L. Global Burden of Alzheimer’s disease and other dementias in adults aged 65 years and older, 1991–2021: population-based study. Front Public Health. Frontiers Media SA; 2025;13. PMID: 40666154

8. Liu H, Nie X, Wang F, Chen D, Zeng Z, Shu P, Huang J. An integrated transcriptomic analysis of brain aging and strategies for healthy aging. Front Aging Neurosci. Frontiers Media SA; 2024;16.

9. Soreq L, Rose J, Soreq E, Hardy J, Trabzuni D, Cookson MR, Smith C, Ryten M, Patani R, Ule J. Major Shifts in Glial Regional Identity Are a Transcriptional Hallmark of Human Brain Aging. Cell Rep. Elsevier B.V.; 2017 Jan 10;18(2):557–570. PMID: 28076797

10. Zhang B, Zhu J. Identification of Key Causal Regulators in Gene Networks. Proceedings of the World Congress on Engineering. 2013;2.

11. Wang M, Roussos P, McKenzie A, Zhou X, Kajiwara Y, Brennand KJ, De Luca GC, Crary JF, Casaccia P, Buxbaum JD, Ehrlich M, Gandy S, Goate A, Katsel P, Schadt E, Haroutunian V, Zhang B. Integrative network analysis of nineteen brain regions identifies molecular signatures and networks underlying selective regional vulnerability to Alzheimer’s disease. Genome Med. 2016;8(1). PMID: 27799057

12. Raj T, Li YI, Wong G, Humphrey J, Wang M, Ramdhani S, Wang YC, Ng B, Gupta I, Haroutunian V, Schadt EE, Young-Pearse T, Mostafavi S, Zhang B, Sklar P, Bennett DA, De Jager PL. Integrative transcriptome analyses of the aging brain implicate altered splicing in Alzheimer’s disease susceptibility. Nat Genet. 2018;

13. Zhang B, Gaiteri C, Bodea LG, Wang Z, McElwee J, Podtelezhnikov AA, Zhang C, Xie T, Tran L, Dobrin R, Fluder E, Clurman B, Melquist S, Narayanan M, Suver C, Shah H, Mahajan M, Gillis T, Mysore J, MacDonald ME, Lamb JR, Bennett DA, Molony C, Stone DJ, Gudnason V, Myers AJ, Schadt EE, Neumann H, Zhu J, Emilsson V. Integrated systems approach identifies genetic nodes and networks in late-onset Alzheimer’s disease. Cell. 2013; PMID: 23622250

14. Wang M, Beckmann ND, Roussos P, Wang E, Zhou X, Wang Q, Ming C, Neff R, Ma W, Fullard JF, Hauberg ME, Bendl J, Peters MA, Logsdon B, Wang P, Mahajan M, Mangravite LM, Dammer EB, Duong DM, Lah JJ, Seyfried NT, Levey AI, Buxbaum JD, Ehrlich M, Gandy S, Katsel P, Haroutunian V, Schadt E, Zhang B. The Mount Sinai cohort of large-scale genomic, transcriptomic and proteomic data in Alzheimer’s disease. Sci Data. England; 2018 Sep;5:180185. PMID: 30204156

15. Wang M, Li A, Sekiya M, Beckmann ND, Quan X, Schrode N, Fernando MB, Yu A, Zhu L, Cao J, Lyu L, Horgusluoglu E, Wang Q, Guo L, Wang YS, Neff R, Song WM, Wang E, Shen Q, Zhou X, Ming C, Ho SM, Vatansever S, Kaniskan HÜ, Jin J, Zhou MM, Ando K, Ho L, Slesinger PA, Yue Z, Zhu J, Katsel P, Gandy S, Ehrlich ME, Fossati V, Noggle S, Cai D, Haroutunian V, Iijima KM, Schadt E, Brennand KJ, Zhang B. Transformative Network Modeling of Multi-omics Data Reveals Detailed Circuits, Key Regulators, and Potential Therapeutics for Alzheimer’s Disease. Neuron. 2021 Jan;109(2):257–272.e14. PMID: 33238137

16. Allen M, Carrasquillo MM, Funk C, Heavner BD, Zou F, Younkin CS, Burgess JD, Chai HS, Crook J, Eddy JA, Li H, Logsdon B, Peters MA, Dang KK, Wang X, Serie D, Wang C, Nguyen T, Lincoln S, Malphrus K, Bisceglio G, Li M, Golde TE, Mangravite LM, Asmann Y, Price ND, Petersen RC, Graff-Radford NR, Dickson DW, Younkin SG, Ertekin-Taner N. Human whole genome genotype and transcriptome data for Alzheimer’s and other neurodegenerative diseases. Sci Data. 2016;3. PMID: 27727239

17. Sun N, Akay LA, Murdock MH, Park Y, Galiana-Melendez F, Bubnys A, Galani K, Mathys H, Jiang X, Ng AP, Bennett DA, Tsai LH, Kellis M. Single-nucleus multiregion transcriptomic analysis of brain vasculature in Alzheimer’s disease. Nat Neurosci. United States; 2023 Jun;26(6):970–982. PMID: 37264161

18. Mathys H, Davila-Velderrain J, Peng Z, Gao F, Mohammadi S, Young JZ, Menon M, He L, Abdurrob F, Jiang X, Martorell AJ, Ransohoff RM, Hafler BP, Bennett DA, Kellis M, Tsai LH. Single-cell transcriptomic analysis of Alzheimer’s disease. Nature. Springer US; 2019;2(May).

19. Mathys H, Boix CA, Akay LA, Xia Z, Davila-Velderrain J, Ng AP, Jiang X, Abdelhady G, Galani K, Mantero J, Band N, James BT, Babu S, Galiana-Melendez F, Louderback K, Prokopenko D, Tanzi RE, Bennett DA, Tsai LH, Kellis M. Single-cell multiregion dissection of Alzheimer’s disease. Nature. England; 2024 Aug;632(8026):858–868. PMID: 39048816

20. Briggs RG, Tanglay O, Dadario NB, Young IM, Fonseka RDi, Hormovas J, Dhanaraj V, Lin YH, Kim SJ, Bouvette A, Chakraborty AR, Milligan TM, Abraham CJ, Anderson CD, O’Donoghue DL, Sughrue ME. The Unique Fiber Anatomy of Middle Temporal Gyrus Default Mode Connectivity. Operative Neurosurgery. Oxford University Press; 2021 Jul 1;21(1):E8–E14. PMID: 33929019

21. Binder JR, Desai RH, Graves WW, Conant LL. Where is the semantic system? A critical review and meta-analysis of 120 functional neuroimaging studies. Cereb Cortex. 2009 Dec;19(12):2767–96. PMID: 19329570

22. Visser M, Jefferies E, Embleton K V, Lambon Ralph MA. Both the middle temporal gyrus and the ventral anterior temporal area are crucial for multimodal semantic processing: distortion-corrected fMRI evidence for a double gradient of information convergence in the temporal lobes. J Cogn Neurosci. 2012 Aug;24(8):1766–78. PMID: 22621260

23. Andrews-Hanna JR. The brain’s default network and its adaptive role in internal mentation. Neuroscientist. 2012 Jun;18(3):251–70. PMID: 21677128

24. Greicius MD, Srivastava G, Reiss AL, Menon V. Default-mode network activity distinguishes Alzheimer’s disease from healthy aging: evidence from functional MRI. Proc Natl Acad Sci U S A. 2004 Mar 30;101(13):4637–42. PMID: 15070770

25. Fjell AM, Westlye LT, Amlien I, Espeseth T, Reinvang I, Raz N, Agartz I, Salat DH, Greve DN, Fischl B, Dale AM, Walhovd KB. High consistency of regional cortical thinning in aging across multiple samples. Cereb Cortex. 2009 Sep;19(9):2001–12. PMID: 19150922

26. Hoffman JM, Welsh-Bohmer KA, Hanson M, Crain B, Hulette C, Earl N, Coleman RE. FDG PET imaging in patients with pathologically verified dementia. J Nucl Med. 2000 Nov;41(11):1920–8. PMID: 11079505

27. Convit A, de Asis J, de Leon MJ, Tarshish CY, De Santi S, Rusinek H. Atrophy of the medial occipitotemporal, inferior, and middle temporal gyri in non-demented elderly predict decline to Alzheimer’s disease. Neurobiol Aging. 2000;21(1):19–26. PMID: 10794844

28. Rogalski EJ, Martersteck A, Roberts AC, Huentelman MJ, Okonkwo OC, Timpo P, Peirce H, Flowers E, Moore S, Zolliecoffer C, Schafer R, Devine R, Engelmeyer J, Geula C, Addison E, Piras I, Trammell AR, Lincoln GA, Goldstein F, Van Ooteghem K, McIlroy B, Bartha R, Finger E, Culum I, Lim A, Swartz RH, Gill NP, Maher AC. The Multisite SuperAging Research Initiative: Enrollment and Scientific Progress. Alzheimer’s & Dementia. Wiley; 2024 Dec;20(S3).

29. Rogalski EJ, Gefen T, Shi J, Samimi M, Bigio E, Weintraub S, Geula C, Mesulam MM. Youthful memory capacity in old brains: anatomic and genetic clues from the Northwestern SuperAging Project. J Cogn Neurosci. 2013 Jan;25(1):29–36. PMID: 23198888

30. Cook Maher A, Makowski-Woidan B, Kuang A, Zhang H, Weintraub S, Mesulam MM, Rogalski E. Neuropsychological Profiles of Older Adults with Superior versus Average Episodic Memory: The Northwestern “SuperAger” Cohort. J Int Neuropsychol Soc. 2022 Jul;28(6):563–573. PMID: 34433508

31. Cook AH, Sridhar J, Ohm D, Rademaker A, Mesulam MM, Weintraub S, Rogalski E. Rates of Cortical Atrophy in Adults 80 Years and Older With Superior vs Average Episodic Memory. JAMA. 2017 Apr 4;317(13):1373–1375. PMID: 28384819

32. Beach TG, Sue LI, Walker DG, Roher AE, Lue L, Vedders L, Connor DJ, Sabbagh MN, Rogers J. The Sun Health Research Institute Brain Donation Program: description and experience, 1987-2007. Cell Tissue Bank. 2008 Sep;9(3):229–45. PMID: 18347928

33. Beach TG, Adler CH, Sue LI, Serrano G, Shill HA, Walker DG, Lue L, Roher AE, Dugger BN, Maarouf C, Birdsill AC, Intorcia A, Saxon-Labelle M, Pullen J, Scroggins A, Filon J, Scott S, Hoffman B, Garcia A, Caviness JN, Hentz JG, Driver-Dunckley E, Jacobson SA, Davis KJ, Belden CM, Long KE, Malek-Ahmadi M, Powell JJ, Gale LD, Nicholson LR, Caselli RJ, Woodruff BK, Rapscak SZ, Ahern GL, Shi J, Burke AD, Reiman EM, Sabbagh MN. Arizona Study of Aging and Neurodegenerative Disorders and Brain and Body Donation Program. Neuropathology. 2015 Aug;35(4):354–89. PMID: 25619230

34. Corrada MM, Berlau DJ, Kawas CH. A population-based clinicopathological study in the oldest-old: the 90+ study. Curr Alzheimer Res. 2012 Jul;9(6):709–17. PMID: 22471863

35. Whittle C, Corrada MM, Dick M, Ziegler R, Kahle-Wrobleski K, Paganini-Hill A, Kawas C. Neuropsychological data in nondemented oldest old: the 90+ Study. J Clin Exp Neuropsychol. 2007 Apr;29(3):290–9. PMID: 17454349

36. Montine TJ, Phelps CH, Beach TG, Bigio EH, Cairns NJ, Dickson DW, Duyckaerts C, Frosch MP, Masliah E, Mirra SS, Nelson PT, Schneider JA, Thal DR, Trojanowski JQ, Vinters H V, Hyman BT, National Institute on Aging, Alzheimer’s Association. National Institute on Aging-Alzheimer’s Association guidelines for the neuropathologic assessment of Alzheimer’s disease: a practical approach. Acta Neuropathol. 2012 Jan;123(1):1–11. PMID: 22101365

37. Hyman BT, Phelps CH, Beach TG, Bigio EH, Cairns NJ, Carrillo MC, Dickson DW, Duyckaerts C, Frosch MP, Masliah E, Mirra SS, Nelson PT, Schneider JA, Thal DR, Thies B, Trojanowski JQ, Vinters H V, Montine TJ. National Institute on Aging-Alzheimer’s Association guidelines for the neuropathologic assessment of Alzheimer’s disease. Alzheimers Dement. 2012 Jan;8(1):1–13. PMID: 22265587

38. Corrada MM, Brookmeyer R, Paganini-Hill A, Berlau D, Kawas CH. Dementia incidence continues to increase with age in the oldest old: the 90+ study. Ann Neurol. 2010 Jan;67(1):114–21. PMID: 20186856

39. Corrada MM, Brookmeyer R, Berlau D, Paganini-Hill A, Kawas CH. Prevalence of dementia after age 90: results from the 90+ study. Neurology. 2008 Jul 29;71(5):337–43. PMID: 18596243

40. Cook Maher A, Makowski-Woidan B, Kuang A, Zhang H, Weintraub S, Mesulam MM, Rogalski E. Neuropsychological Profiles of Older Adults with Superior versus Average Episodic Memory: The Northwestern “SuperAger” Cohort. J Int Neuropsychol Soc. 2022 Jul;28(6):563–573. PMID: 34433508

41. Rogalski EJ, Gefen T, Shi J, Samimi M, Bigio E, Weintraub S, Geula C, Mesulam MM. Youthful memory capacity in old brains: anatomic and genetic clues from the Northwestern SuperAging Project. J Cogn Neurosci. 2013 Jan;25(1):29–36. PMID: 23198888

42. Pletnikova O, Rudow GL, Hyde TM, Kleinman JE, Ali SZ, Bharadwaj R, Gangadeen S, Crain BJ, Fowler DR, Rubio AI, Troncoso JC. Alzheimer Lesions in the Autopsied Brains of People 30 to 50 Years of Age. Cogn Behav Neurol. 2015 Sep;28(3):144–52. PMID: 26413742

43. Pletnikova O, Kageyama Y, Rudow G, LaClair KD, Albert M, Crain BJ, Tian J, Fowler D, Troncoso JC. The spectrum of preclinical Alzheimer’s disease pathology and its modulation by ApoE genotype. Neurobiol Aging. 2018 Nov;71:72–80. PMID: 30099348

44. Dobin A, Davis CA, Schlesinger F, Drenkow J, Zaleski C, Jha S, Batut P, Chaisson M, Gingeras TR. STAR: Ultrafast universal RNA-seq aligner. Bioinformatics. 2013;29(1):15–21. PMID: 23104886

45. Okonechnikov K, Conesa A, García-Alcalde F. Qualimap 2: Advanced multi-sample quality control for high-throughput sequencing data. Bioinformatics. 2015;32(2):292–294. PMID: 26428292

46. Love MI, Huber W, Anders S. Moderated estimation of fold change and dispersion for RNA-seq data with DESeq2. Genome Biol [Internet]. 2014;15(12):550. Available from: http://genomebiology.biomedcentral.com/articles/10.1186/s13059-014-0550-8 PMID: 25516281

47. Benjamini Y, Hochberg Y. Benjamini Y, Hochberg Y. Controlling the false discovery rate: a practical and powerful approach to multiple testing. Journal of the Royal Statistical Society B [Internet]. 1995;57(1):289–300. Available from: http://www.stat.purdue.edu/~doerge/BIOINFORM.D/FALL06/Benjamini and YFDR.pdf%5Cnhttp://engr.case.edu/ray_soumya/mlrg/controlling_fdr_benjamini95.pdf PMID: 11682119

48. Zhu A, Ibrahim JG, Love MI. Heavy-tailed prior distributions for sequence count data: removing the noise and preserving large differences. Bioinformatics. 2019 Jun 1;35(12):2084–2092. PMID: 30395178

49. Ritchie ME, Phipson B, Wu D, Hu Y, Law CW, Shi W, Smyth GK. limma powers differential expression analyses for RNA-sequencing and microarray studies. Nucleic Acids Res. 2015;43(7):e47. PMID: 25605792

50. Yu G, Wang LG, Han Y, He QY. clusterProfiler: an R package for comparing biological themes among gene clusters. OMICS. 2012 May;16(5):284–287. PMID: 22455463

51. Piras IS, Huentelman MJ, Pinna F, Paribello P, Solmi M, Murru A, Carpiniello B, Manchia M, Zai CC. A review and meta-analysis of gene expression profiles in suicide. Eur Neuropsychopharmacol. Netherlands; 2021 Dec;56:39–49. PMID: 34923210

52. Korotkevich G, Sukhov V. Fast gene set enrichment analysis. 2016;1–29.

53. Song WM, Zhang B. Multiscale Embedded Gene Co-expression Network Analysis. PLoS Comput Biol. 2015 Nov;11(11):e1004574. PMID: 26618778

54. Langfelder P, Horvath S. WGCNA: an R package for weighted correlation network analysis. BMC Bioinformatics. 2008;9:559. PMID: 19114008

55. Shu L, Zhao Y, Kurt Z, Byars SG, Tukiainen T, Kettunen J, Orozco LD, Pellegrini M, Lusis AJ, Ripatti S, Zhang B, Inouye M, Mäkinen VP, Yang X. Mergeomics: multidimensional data integration to identify pathogenic perturbations to biological systems. BMC Genomics. England; 2016 Nov;17(1):874. PMID: 27814671

56. Brandenberger R, Schmidt A, Linton J, Wang D, Backus C, Denda S, Müller U, Reichardt LF. Identification and characterization of a novel extracellular matrix protein nephronectin that is associated with integrin α8β1 in the embryonic kidney. Journal of Cell Biology. 2001 Jul 23;154(2):447–458. PMID: 11470831

57. Felsky D, Santa-Maria I, Cosacak I, French L, Schneider JA, Bennett DA, De Jager PL, Kizil C, Tosto G. The Caribbean-Hispanic Alzheimer’s disease brain transcriptome reveals ancestry-specific disease mechanisms. Available from: 10.7303/syn2580853

58. McCorkindale AN, Patrick E, Duce JA, Guennewig B, Sutherland GT. The Key Factors Predicting Dementia in Individuals With Alzheimer’s Disease-Type Pathology. Front Aging Neurosci. Frontiers Media S.A.; 2022 Apr 25;14.

59. Logue MW, Labadorf A, O’Neill NK, Dickson DW, Dugger BN, Flanagan ME, Frosch MP, Gearing M, Jin LW, Kofler J, Mayeux R, McKee A, Miller CA, Murray ME, Nelson PT, Perrin RJ, Schneider JA, Stein TD, Teich AF, Troncoso JC, Wang SH, Wolozin B, Mez J, Farrer LA. Transcriptome-wide association study of Alzheimer disease reveals many differentially expressed genes and multiple biological pathways in brain tissue from African American donors. 2024. Available from: http://medrxiv.org/lookup/doi/10.1101/2024.10.29.24316311

60. Ýþ Ö, Wang X, Reddy JS, Min Y, Yilmaz E, Bhattarai P, Patel T, Bergman J, Quicksall Z, Heckman MG, Tutor-New FQ, Can Demirdogen B, White L, Koga S, Krause V, Inoue Y, Kanekiyo T, Cosacak MI, Nelson N, Lee AJ, Vardarajan B, Mayeux R, Kouri N, Deniz K, Carnwath T, Oatman SR, Lewis-Tuffin LJ, Nguyen T, Carrasquillo MM, Graff-Radford J, Petersen RC, Jack CR, Kantarci K, Murray ME, Nho K, Saykin AJ, Dickson DW, Kizil C, Allen M, Ertekin-Taner N. Gliovascular transcriptional perturbations in Alzheimer’s disease reveal molecular mechanisms of blood brain barrier dysfunction. Nat Commun. Nature Research; 2024 Dec 1;15(1). PMID: 38902234

61. Yang AC, Vest RT, Kern F, Lee DP, Agam M, Maat CA, Losada PM, Chen MB, Schaum N, Khoury N, Toland A, Calcuttawala K, Shin H, Pálovics R, Shin A, Wang EY, Luo J, Gate D, Schulz-Schaeffer WJ, Chu P, Siegenthaler JA, McNerney MW, Keller A, Wyss-Coray T. A human brain vascular atlas reveals diverse mediators of Alzheimer’s risk. Nature. Nature Research; 2022 Mar 31;603(7903):885–892. PMID: 35165441

62. Wan YW, Al-Ouran R, Mangleburg CG, Perumal TM, Lee T V., Allison K, Swarup V, Funk CC, Gaiteri C, Allen M, Wang M, Neuner SM, Kaczorowski CC, Philip VM, Howell GR, Martini-Stoica H, Zheng H, Mei H, Zhong X, Kim JW, Dawson VL, Dawson TM, Pao PC, Tsai LH, Haure-Mirande JV, Ehrlich ME, Chakrabarty P, Levites Y, Wang X, Dammer EB, Srivastava G, Mukherjee S, Sieberts SK, Omberg L, Dang KD, Eddy JA, Snyder P, Chae Y, Amberkar S, Wei W, Hide W, Preuss C, Ergun A, Ebert PJ, Airey DC, Mostafavi S, Yu L, Klein HU, Carter GW, Collier DA, Golde TE, Levey AI, Bennett DA, Estrada K, Townsend TM, Zhang B, Schadt E, De Jager PL, Price ND, Ertekin-Taner N, Liu Z, Shulman JM, Mangravite LM, Logsdon BA. Meta-Analysis of the Alzheimer’s Disease Human Brain Transcriptome and Functional Dissection in Mouse Models. Cell Rep. Elsevier B.V.; 2020 Jul 14;32(2).

63. Nguyen PT, Dorman LC, Pan S, Vainchtein ID, Han RT, Nakao-Inoue H, Taloma SE, Barron JJ, Molofsky AB, Kheirbek MA, Molofsky A V. Microglial Remodeling of the Extracellular Matrix Promotes Synapse Plasticity. Cell. 2020 Jul 23;182(2):388–403.e15. PMID: 32615087

64. Soto JS, Jami-Alahmadi Y, Chacon J, Moye SL, Diaz-Castro B, Wohlschlegel JA, Khakh BS. Astrocyte-neuron subproteomes and obsessive-compulsive disorder mechanisms. Nature. 2023 Apr;616(7958):764–773. PMID: 37046092

65. Hill MA, Gammie SC. Alzheimer’s disease large-scale gene expression portrait identifies exercise as the top theoretical treatment. Sci Rep. Nature Research; 2022 Dec 1;12(1). PMID: 36229643

66. Zhong L, Sheng X, Wang W, Li Y, Zhuo R, Wang K, Zhang L, Hu DD, Hong Y, Chen L, Rao H, Li T, Chen M, Lin Z, Zhang Y wu, Wang X, Yan XX, Chen X, Bu G, Chen XF. TREM2 receptor protects against complement-mediated synaptic loss by binding to complement C1q during neurodegeneration. Immunity. Cell Press; 2023 Aug 8;56(8):1794–1808.e8. PMID: 37442133

67. Dejanovic B, Wu T, Tsai MC, Graykowski D, Gandham VD, Rose CM, Bakalarski CE, Ngu H, Wang Y, Pandey S, Rezzonico MG, Friedman BA, Edmonds R, De Mazière A, Rakosi-Schmidt R, Singh T, Klumperman J, Foreman O, Chang MC, Xie L, Sheng M, Hanson JE. Complement C1q-dependent excitatory and inhibitory synapse elimination by astrocytes and microglia in Alzheimer’s disease mouse models. Nat Aging. Springer; 2022 Sep 1;2(9):837–850.

68. Valiukas Z, Tangalakis K, Apostolopoulos V, Feehan J. Microglial activation states and their implications for Alzheimer’s Disease. The journal of prevention of Alzheimer’s disease. 2025. p. 100013. PMID: 39800461

69. Guo F, Sheng ZH, Fu Y, Wang ZB, Xue RJ, Tan L, Tan MS, Wang ZT. Complement C1q is associated with neuroinflammation and mediates the association between amyloid-β and tau pathology in Alzheimer’s disease. Transl Psychiatry. Springer Nature; 2025 Dec 1;15(1). PMID: 40675947

70. Mukherjee S, Klaus C, Pricop-Jeckstadt M, Miller JA, Struebing FL. A microglial signature directing human aging and neurodegeneration-related gene networks. Front Neurosci. Frontiers Media S.A.; 2019;13(JAN).

71. Buncha V, Fopiano KA, Lang L, Williams C, Horuzsko A, Filosa JA, Kapuku G, Bagi Z. Mice with endothelial cell-selective adhesion molecule deficiency develop coronary microvascular rarefaction and left ventricle diastolic dysfunction. Physiol Rep. American Physiological Society; 2023 Mar 1;11(6). PMID: 36946064

72. Mostafavi S, Gaiteri C, Sullivan SE, White CC, Tasaki S, Xu J, Taga M, Klein HU, Patrick E, Komashko V, McCabe C, Smith R, Bradshaw EM, Root DE, Regev A, Yu L, Chibnik LB, Schneider JA, Young-Pearse TL, Bennett DA, De Jager PL. A molecular network of the aging human brain provides insights into the pathology and cognitive decline of Alzheimer’s disease. Nat Neurosci. 2018;

73. Kaushansky N, Eisenstein M, Zilkha-Falb R, Ben-Nun A. The myelin-associated oligodendrocytic basic protein (MOBP) as a relevant primary target autoantigen in multiple sclerosis. Autoimmunity Reviews. 2010. p. 233–236. PMID: 19683076

74. Bettencourt C, Miki Y, Piras IS, de Silva R, Foti SC, Talboom JS, Revesz T, Lashley T, Balazs R, Viré E, Warner TT, Huentelman MJ, Holton JL. MOBP and HIP1 in multiple system atrophy: New α-synuclein partners in glial cytoplasmic inclusions implicated in the disease pathogenesis. Neuropathol Appl Neurobiol. 2021 Aug;47(5):640–652. PMID: 33368549

75. Sanguinetti C, Minniti M, Susini V, Caponi L, Panichella G, Castiglione V, Aimo A, Emdin M, Vergaro G, Franzini M. The Journey of Human Transthyretin: Synthesis, Structure Stability, and Catabolism. Biomedicines. MDPI; 2022.

76. Greve AM, Christoffersen M, Frikke-Schmidt R, Nordestgaard BG, Tybjærg-Hansen A. Association of Low Plasma Transthyretin Concentration With Risk of Heart Failure in the General Population. JAMA Cardiol. 2021 Mar 1;6(3):258–266. PMID: 33237279

