## Supplementary_figures for "Transcriptomic profiling of the middle temporal gyrus reveals differential glial/neuronal dysregulation across Alzheimer’s disease and aging"

***Figure S1****. Total sequenced reads distribution (in million) in the 621 sequenced samples.*

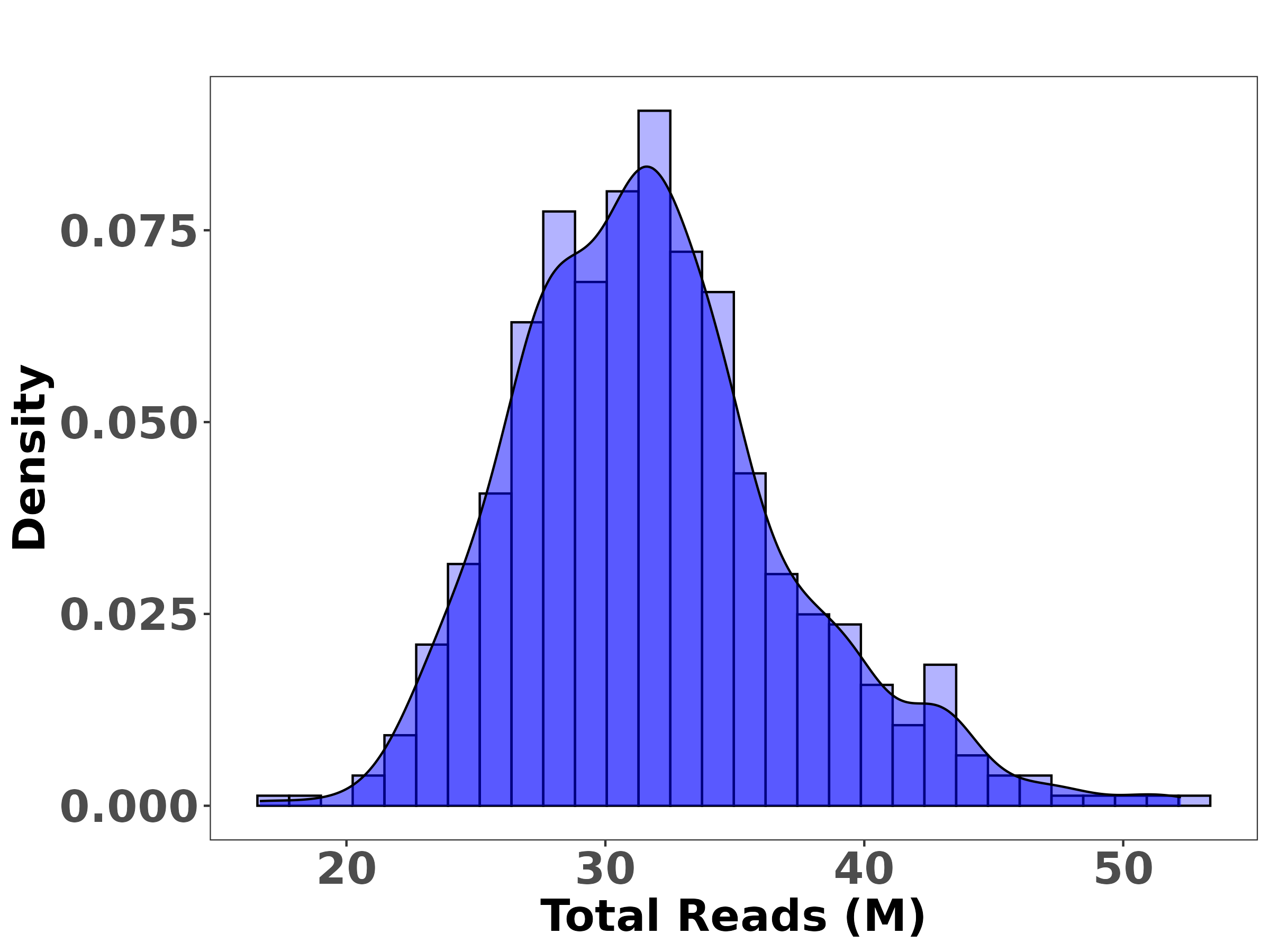

***Figures S2****. Uniquely mapped reads distribution as estimated by STAR in the 621 sequenced samples.*

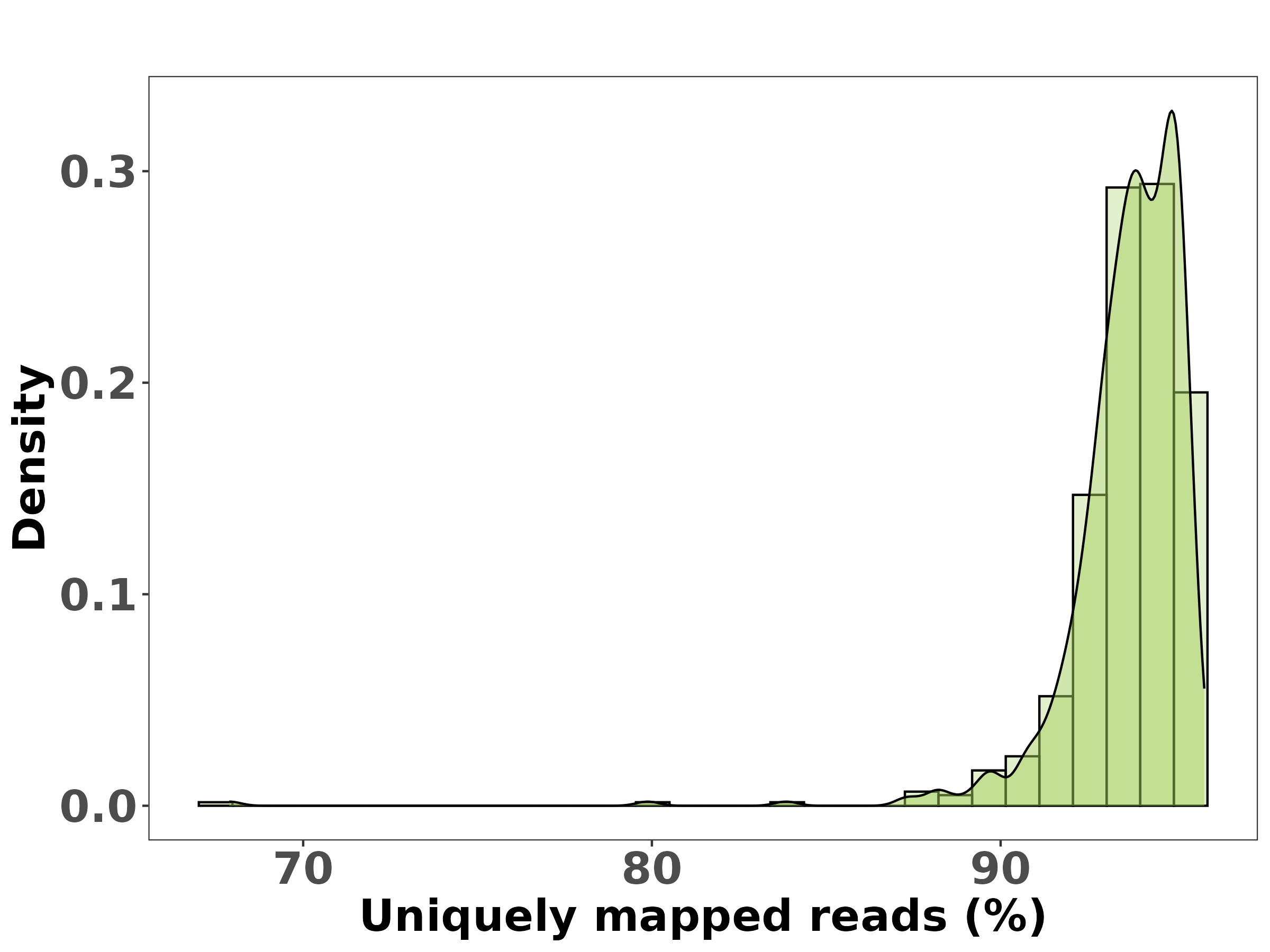

***Fig. S3.*** *Plot of the two top principal components. Red points are the 11 identified outliers (± 3 standard deviations across the mean of one of the two top components).*

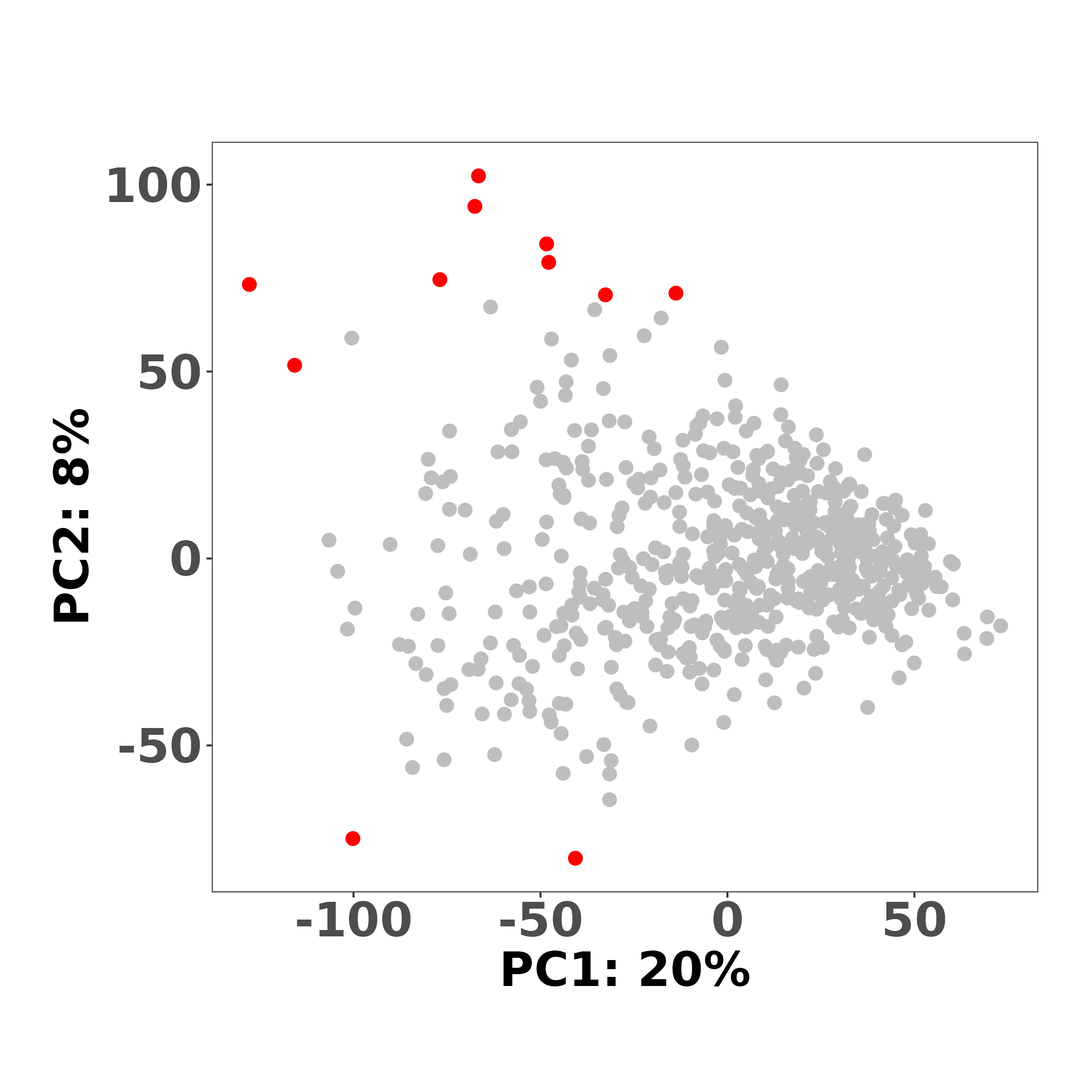

***Fig. S4.*** *Plot of the two top principal components after removing the 11 outliers (final sample size: n = 606).*

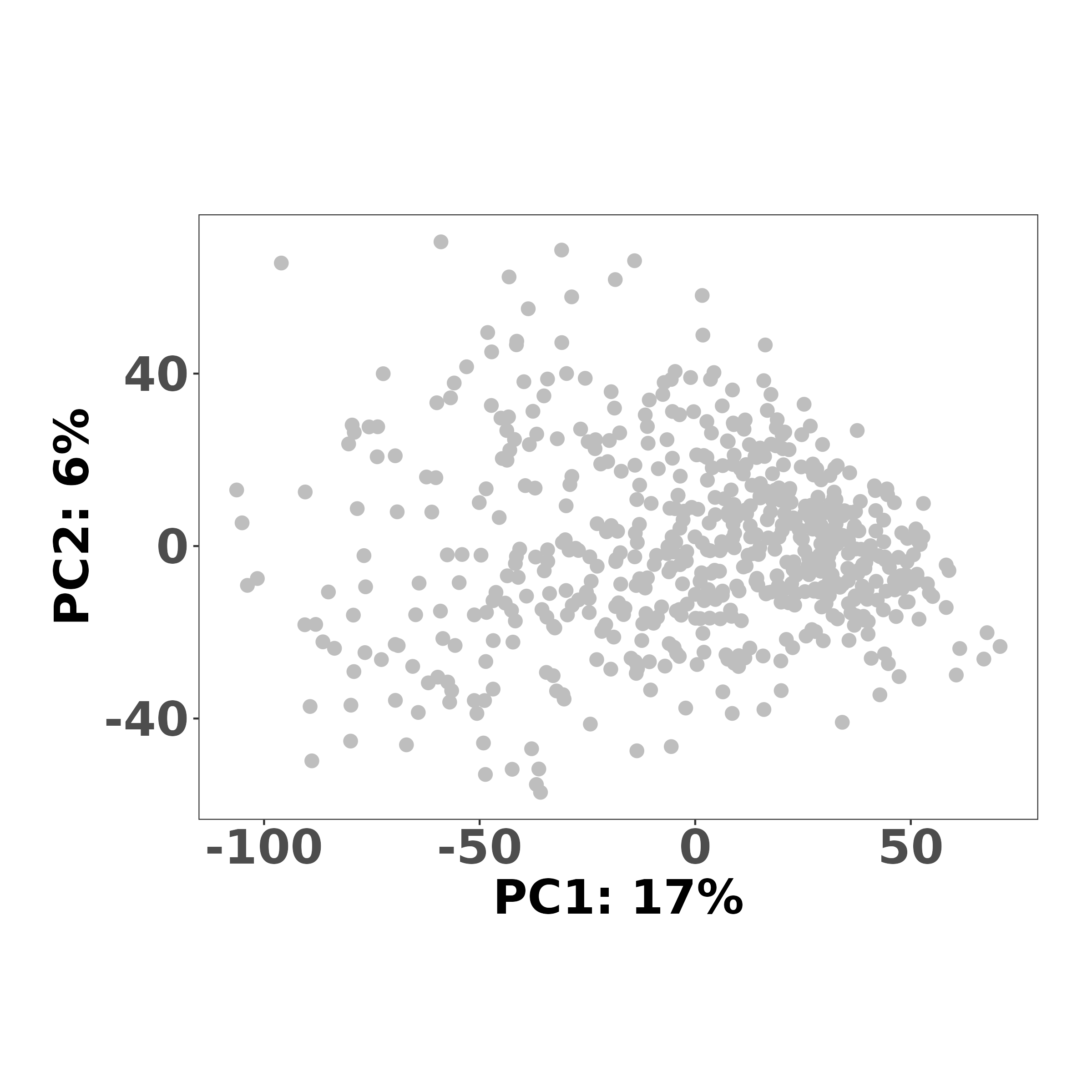
